# Gamma synchronization between V1 and V4 improves behavioral performance

**DOI:** 10.1101/290817

**Authors:** Gustavo Rohenkohl, Conrado Arturo Bosman, Pascal Fries

**Author notes:** Correspondence should be addressed to PF.

## Abstract

Motor behavior is often driven by visual stimuli, relying on efficient feedforward communication from lower to higher visual areas. The Communication-through-Coherence hypothesis proposes that interareal communication depends on coherence at an optimal phase relation. While previous studies have linked effective communication to enhanced interareal coherence, it remains unclear, whether this interareal coherence occurs at an optimal phase relation that actually improves the stimulus transmission to behavioral report. We recorded local field potentials simultaneously from areas V1 and V4 of macaque monkeys performing a selective visual attention task, during which they reported changes of the attended stimulus. Gamma synchronization between V1 and V4, immediately preceding the stimulus change, predicted subsequent reaction times (RTs). Crucially, RTs were systematically slowed as trial-by-trial interareal gamma phase relations deviated from the phase relation at which V1 and V4 synchronized on average. These effects were specific to the attended stimulus and not due to local power or phase inside V1 or V4. We conclude that interareal gamma synchronization occurs at the optimal phase relation and thereby improves interareal communication and the effective transformation of sensory inputs into motor responses.

## INTRODUCTION

At the heart of many cognitive functions is the dynamic modulation of effective connectivity, i.e. the context-dependent modulation of the postsynaptic impact of a given neuronal group. The impact of a group of neurons can be enhanced when they engage in gamma-band synchronization, because this renders their inputs to postsynaptic target neurons coincident in time (Azouz and Gray, 2003; Salinas and Sejnowski, 2000). Indeed, groups of visual cortical neurons show enhanced local gamma-band synchronization when they process attended as compared to ignored stimuli (Bichot et al., 2005; Fries et al., 2001; Taylor et al., 2005), and this enhancement predicts reaction times (Womelsdorf et al., 2006). Thus, effective connectivity depends on synchronization among presynaptic neurons.

Importantly, effective connectivity also depends on the synchronization between pre- and postsynaptic neurons. When postsynaptic neurons are synchronized, this modulates their gain rhythmically, and inputs are most effective when they consistently arrive during high gain. Thus, good neuronal communication requires coherence between pre- and postsynaptic neurons, a concept referred to as Communication-trough-Coherence (CTC) (Akam and Kullmann, 2010; Börgers and Kopell, 2008; Fries, 2015; Palmigiano et al., 2017). Indeed, gamma phase relations between neuronal groups affect their power-power correlation (Womelsdorf et al., 2007) and their transfer entropy (Besserve et al., 2015). Furthermore, this mechanism might actually serve a functional role for selective attention. When two visual stimuli induce two local gamma rhythms in macaque V1, only the gamma rhythm induced by the attended stimulus establishes coherence to V4 (Bosman et al., 2012; Grothe et al., 2012).

Whether interareal coherence can in fact affect interareal communication according to the CTC mechanism, depends on whether the postsynaptic gamma rhythm modulates the gain of spike responses. One study investigated whether a spike in V1 is followed by a spike in V2, and found this spike transmission modulated by the gamma phase in V2 (Jia et al., 2013). Another recent study showed that visually induced gamma in V4 rhythmically modulates the gain of spike responses and also behavioral reaction times (Ni et al., 2016). Similar effects have been described for gamma-frequency rhythms that were optogenetically entrained in rodent somatosensory cortex (Cardin et al., 2009; Siegle et al., 2014). Thus, several studies have shown that postsynaptic gamma rhythmically modulates gain.

However, it remains to be shown that pre- and post-synaptic neurons engage in coherence at the phase relation that actually improves communication. The studies on selective attention effects report enhancements of coherence. But enhanced coherence merely reflects an enhanced consistency of phase relations. A consistent phase relation is required for good communication, because it is required to consistently time inputs to postsynaptic phases of high gain. Yet, a consistent phase relation is not sufficient for good communication, because it can as well consistently time inputs to postsynaptic phases of low gain. Thus, while coherence is necessary for communication, it is not sufficient, and the other crucial factor is the phase relation at which pre- and post-synaptic neurons are coherent (Akam and Kullmann, 2010; Fries, 2005). Similarly, the studies showing gamma-rhythmic gain modulation do not show that presynaptic activity is coherent at such a phase relation that inputs are timed to moments of maximal postsynaptic gain. Thus, one of the most important requirements of CTC, namely that coherence occurs at the optimal phase relation for communication, has yet to receive empirical support. Therefore, we set out to test whether interareal gamma synchronization actually occurs at the optimal interareal phase relation and thereby improves behavioral performance.

To address these issues directly, we investigated simultaneous V1-V4 recordings in macaques performing a selective visual attention task to test whether the transmission of a stimulus change to a behavioral response depends on the V1-V4 phase relation. As the stimulus change is timed randomly, it is independent of ongoing neuronal activity and provides an ideal probe for the efficiency of transmission. Furthermore, the analysis relates interareal synchronization directly to behavioral responses, thereby directly probing its behavioral relevance. We first describe that fast responses are preceded by enhanced V1-V4 gamma-band synchronization. We then show that interareal gamma phase relations just before the stimulus change predict the speed, at which the change is transformed into a behavioral response. Most importantly, we show that the phase relation at which V1 and V4 synchronize, is optimal for stimulus transmission: The two areas synchronize at their preferred or mean phase relation, and we show that any deviation from this mean phase relation on a given trial systematically slows behavioral response.

## RESULTS

We intended to investigate the relation between interareal synchronization and behavior. To this end, we focused on neuronal activity preceding the behaviorally relevant stimulus event. The relevant stimulus event was a shape change of a cued drifting grating. The cued and an un-cued grating were simultaneously presented in opposite hemifields, while monkeys kept fixation and attentionally monitored the cued grating (Fig. 1). Trials with attention to the stimulus contralateral to the ECoG are referred to as “attend-IN”, and trials with attention ipsilateral to the ECoG as “attend-OUT”. Unless otherwise noted, the analysis focuses on the 200 ms immediately preceding the change of the attended stimulus and uses the corresponding reaction time (RT).

**Figure 1.**
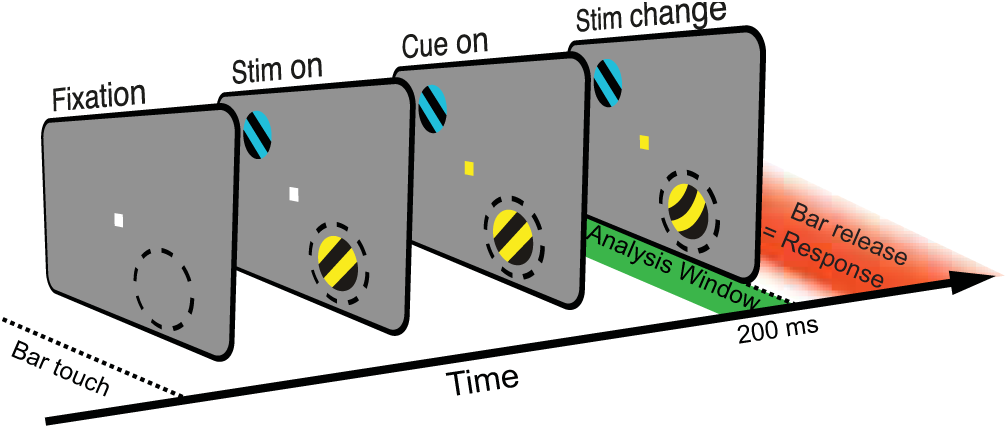
Behavioral task. Macaques were trained to touch a bar, which triggered the appearance of a central fixation point. Two stimuli were presented, one in the RF of the recording sites (illustrated as dashed circle, not visible to the monkey) and one in the opposite quadrant. Blue and yellow tints were randomly assigned to the two stimuli. The fixation point assumed the color of one of the stimuli, cueing that stimulus as the behaviorally relevant, i.e. attended, one. Either one of the stimuli could undergo a bend at an unpredictable time. If the change occurred in the attended stimulus, and the bar was released within a short time window thereafter (illustrated in red), a reward was given. The analysis focused on a 200 ms time window immediately before the stimulus change (illustrated in green). Full details of the task are described in the Methods section.

### Visually induced gamma-band activities in V1 and V4 and their interareal synchronization

During stimulus presentation and task performance, we recorded neuronal activity simultaneously from areas V1 and V4. Both areas were covered with an electrocorticographic (ECoG) grid. Visual stimulation induced gamma band activity in retinotopically corresponding parts of V1 and V4 (Fig. 2a,b)(Lewis et al., 2016b). For further analysis, we selected per monkey the visually driven recording sites, defined as the top third of sites with the strongest visually induced gamma-band activity (see Methods for details). The spectra of stimulation-induced power changes showed clear gamma-band peaks for both V1 (Fig. 2c,d) and V4 (Fig. 2e,f), with particularly strong power increases in V1. The gamma rhythms in V1 and V4 showed interareal synchronization, as evidenced by the spectra of pairwise phase consistency (PPC, Fig. 2g,h; in this and the following analyses of interareal coherence, N=140 interareal site pairs and N=2550 trials). For further analysis, we determined the individual gamma peak frequency for each monkey separately (see Methods for details). Spectra of local power changes and of interareal coherence agreed largely but not perfectly in peak frequency, as has been observed in previous studies of interareal synchronization (Bosman et al., 2012; Gregoriou et al., 2009). As our focus was on determining whether interareal synchronization is predictive of behavior, we determined the individual gamma peak frequency by fitting a Gaussian to the PPC spectrum. The gamma peak frequency was 74 Hz in monkey K and 63 Hz in monkey P. This is in agreement with previous studies showing inter-individual variability in gamma peak frequency (van Pelt et al., 2012; Vinck et al., 2010a).

**Figure 2.**
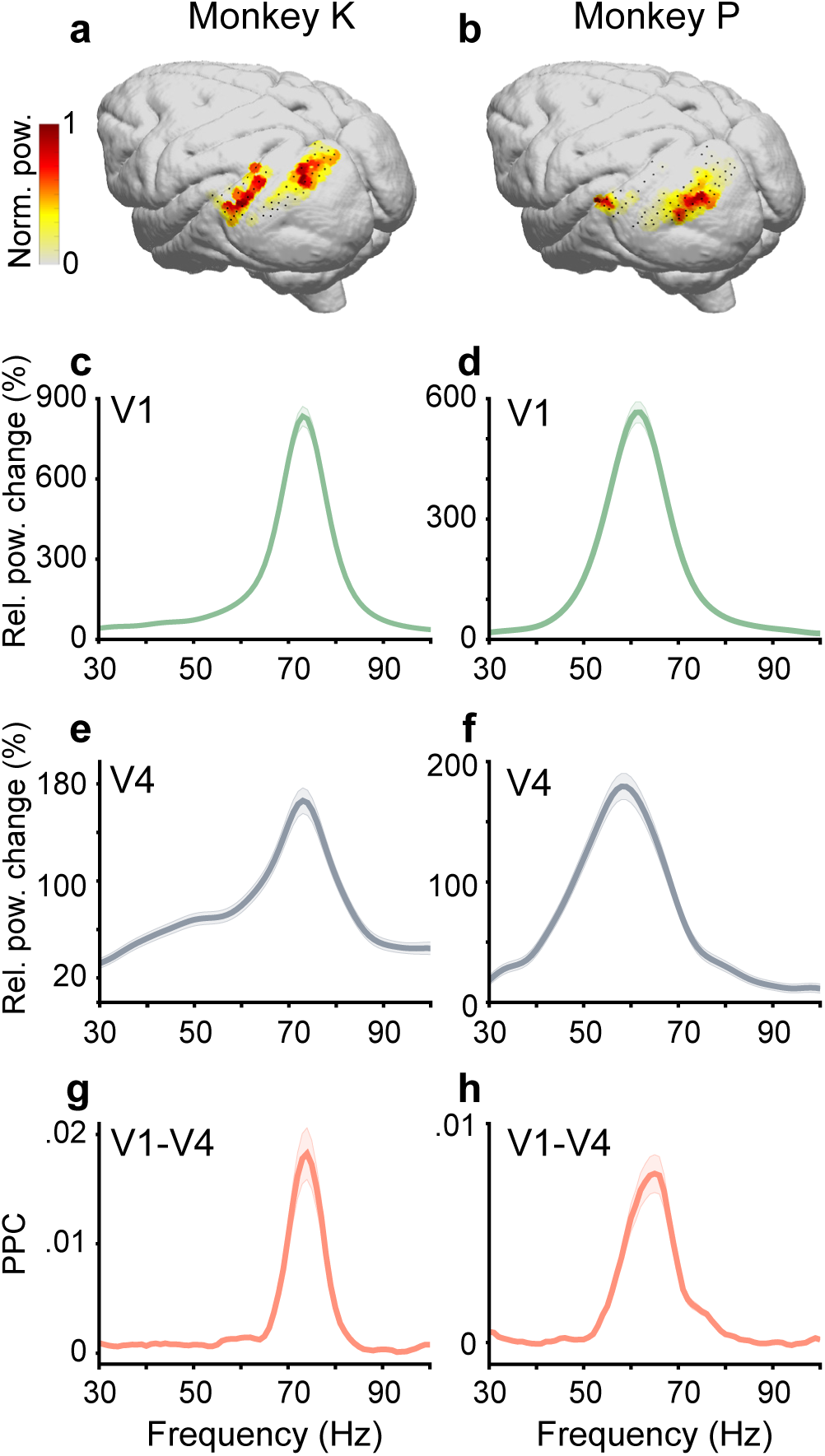
ECoG coverage of areas V1 and V4, and spectra of visually induced power changes and of interareal synchronization. (a) Dots represent positions of ECoG recording sites in monkey K, projected onto a standard brain. Overlaid color map illustrates visually induced gamma-band activity in areas V1 and V4. Note that the color range is scaled separately per area, because visually induced gamma was substantially stronger in V1. (b) Same as a, but for monkey P. (c) Spectrum of visually induced power changes in V1, quantified as percent change relative to pre-stimulus baseline. (d) Same as c, but for monkey P. (e,f) Same as c,d, but for area V4. (g) Spectrum of V1-V4 synchronization in monkey K, as quantified by the pairwise phase consistency (PPC). (h) Same as g, but for monkey P. In c-f, for area V1 and area V4 separately, data were averaged over the top third of recording sites with the strongest visually induced gamma-band activity, because those sites were used for further analysis.

### Short reaction times are preceded by strong interareal gamma-band synchronization

As a first approach to test for a putative relation between interareal gamma-band synchronization and behavior, we performed a median split of the trials according to RTs. Note that RTs did not differ between attend-IN and attend-OUT (p=0.18, Supplementary Fig. S1). For fast and slow trials separately, we calculated the V1-V4 PPC in a 200 ms window immediately preceding the stimulus change. To combine results from both monkeys, PPC spectra were aligned to the individual gamma peak frequencies. V1-V4 PPC was stronger during the attend-IN as compared to the attend-OUT condition (compare Fig. 3a,b; p < 0.05, 2×2 ANOVA with factors Attend-IN/Attend-OUT and RT-fast/RT-slow). This is consistent with previous findings of increased interareal coherence with attention (Bosman et al., 2012; Gregoriou et al., 2009; Grothe et al., 2012; Richter et al., 2017). Importantly, V1 V4 gamma PPC was enhanced before stimulus changes reported with short RTs as compared to long RTs (Fig. 3a, p<0.05, non-parametric randomization test with correction for multiple comparisons across frequencies). This effect was absent for the attend-OUT condition (Fig. 3b), when monkeys reported changes in the ipsilateral stimulus, and thereby the observed RTs related to the stimulus not processed by the recorded neurons. Thus, the RT-related effect of gamma synchronization is specific to the contralateral V1 V4 area pair that is actually involved in the communication of the behaviorally relevant stimulus, and the effect is not due to fluctuations in overall arousal.

**Figure 3.**
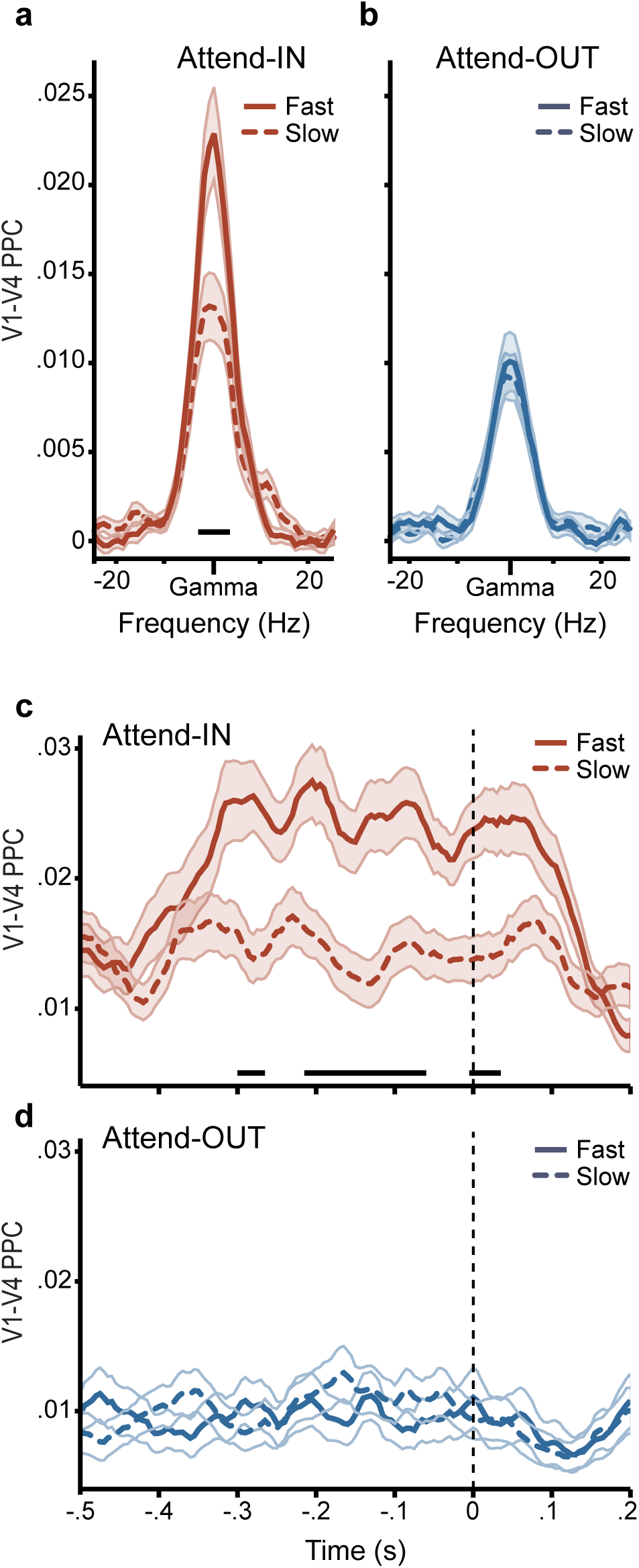
Short reaction times (RTs) are preceded by strong interareal gamma-band synchronization. (a) V1 V4 PPC during attend-IN as a function of frequency relative to the individual gamma peak frequency, calculated for a 200 ms window immediately preceding the onset of stimulus change. PPC spectra from trials with short (long) RTs are shown as solid (dashed) lines. (b) Same as a, but for attend OUT. (c) V1-V4 PPC during attend-IN at the individual gamma peak frequency, as a function of time relative to the onset of stimulus change (vertical dashed line). (d) Same as c, but for attend OUT. Black horizontal bars indicate statistical significance (p<0.05) obtained with a non-parametric randomization test including correction for the multiple comparisons.

The RT-related effect was furthermore specific in time to the period just before and around the stimulus change. The time-resolved PPC analysis during attend-IN reveals a start of the effect around 300 ms before the stimulus change and an end around 100 ms after the stimulus change (Fig. 3c). The absence of an effect during attend-OUT was confirmed in the time-resolved analysis (Fig. 3d).

### Interareal gamma-band phase relations predict reaction times

To test whether interareal synchronization is predictive of behavior, we determined the phase relation immediately preceding the stimulus change and investigated whether it systematically affected the subsequent behavioral reaction times to the stimulus change. As the previous analysis revealed an RT related effect specifically in the gamma band and for attend-IN, we first focused on that band and those trials. We calculated the V1 V4 gamma phase relation for the 200 ms window immediately preceding each stimulus change. Based on those phase relations, we sorted trials into 36 bins and averaged RTs per bin. Figure 4a shows the result of this analysis for an example site pair during attend IN, and suggests a systematic dependence. By contrast, Figure 4b shows the analysis for the same pair during attend-OUT, and suggests a much weaker or no effect. To quantify the effect, we calculated the circular-linear correlation (Berens, 2009) between interareal phase relations and RTs, across trials and without binning, thereby reflecting the true trial-by-trial correlation coefficient (Richter et al., 2015, 2017). The red line in Figure 4c shows the average correlation coefficient over all interareal site pairs and both monkeys during attend-IN as a function of frequency, and reveals a significant correlation in the gamma-band range (p<0.05, non-parametric randomization test corrected for multiple comparisons across frequencies). When we repeated the same analysis for the attend-OUT condition, the correlation was absent.

**Figure 4.**
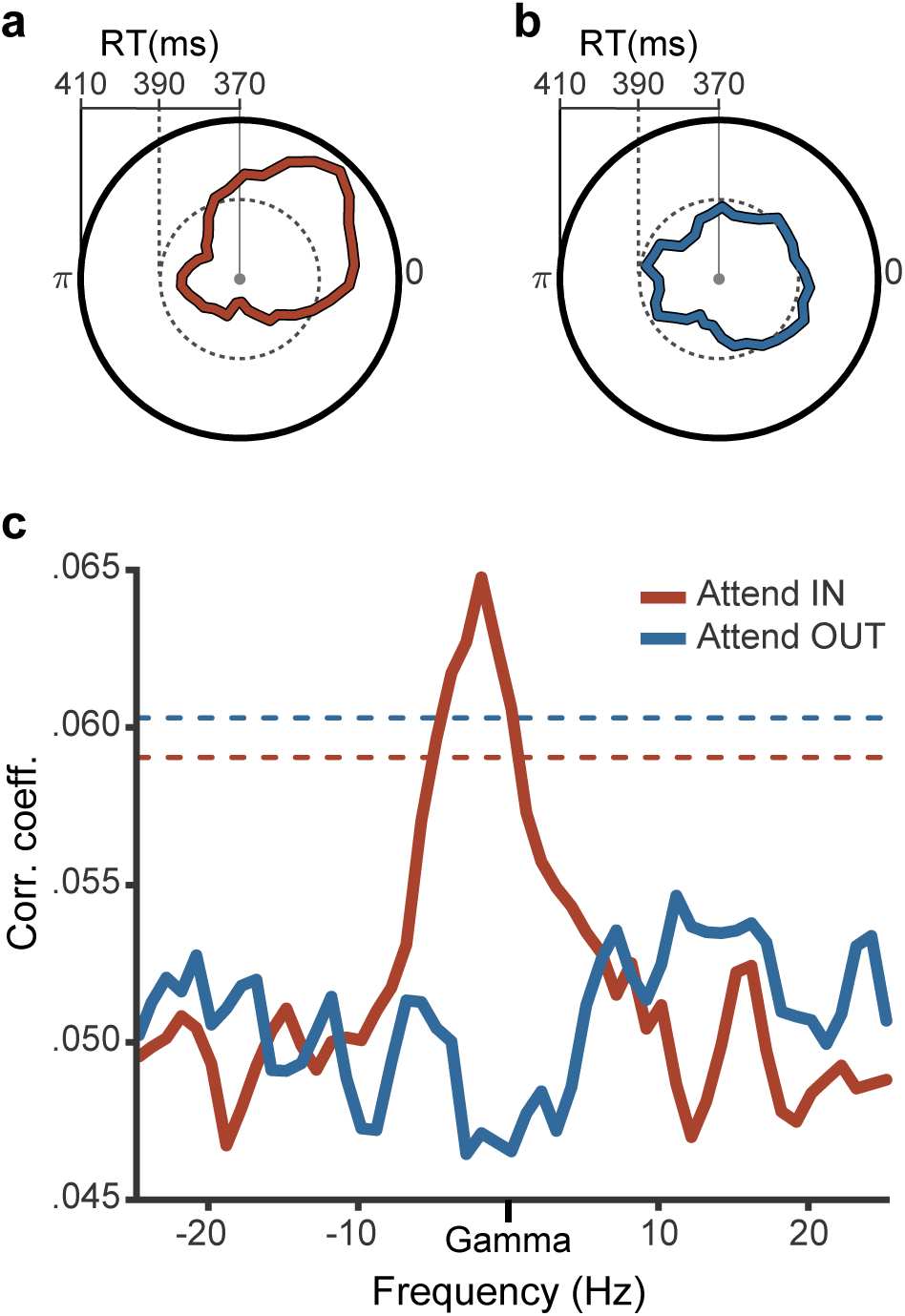
Interareal gamma-band phase relations predict reaction times. (a) Polar plot showing reaction time as a function of V1-V4 gamma phase relation during attend-IN, for one example pair of recording sites. (b) Same as a, but during attend-OUT. (c) Circular-linear correlation between gamma phase relation and RT, averaged over all V1-V4 pairs of recording sites. Data from attend-IN (attend-OUT) are shown in red (blue). The dashed lines indicate the significance thresholds (p<0.05) obtained with a non-parametric randomization test including correction for multiple comparisons.

### Trial-by-trial deviation from mean interareal gamma-band phase relation predicts reaction time

The correlation analysis captures any linear correlation between the phase-relation and RT, irrespective of the actual phase relations resulting in minimal and maximal RTs. Thus, the significant correlation reveals that gamma phase relations are predictive of RTs in the average across interareal site pairs, but the phase relations leading to minimal RTs could differ across site pairs, and even if they were consistent, the phase relation leading to minimal RTs could take some arbitrary and hard-to-interpret value. Specifically, this analysis does not yet establish that the phase relation at which synchronization occurs is optimal for stimulus transmission. Correspondingly, as a next step, we aimed at testing the specific hypothesis that the phase-relation, at which interareal site pairs synchronized, resulted in shortest RTs. Synchronization entails that site pairs spend relatively more time in a particular phase relation (Lowet et al., 2017). Therefore, the phase-relation of synchronization is the mean phase relation. Thus, we hypothesized that the mean phase relation leads to the shortest RTs, and that deviations from this mean phase relation lead to longer RTs. Note that the raw phase relation between a given V1-V4 site pair cannot be directly interpreted, because the absolute LFP phase depends on numerous accidental factors like the geometric relationship between source and electrode, and additionally the bipolar derivation used to remove the common recording reference incurs arbitrary phase rotations. Yet, irrespective of this, the mean phase relation reflects the phase relation of synchronization, and the phase relation in each trial can be expressed in terms of its deviation from this mean phase relation.

Figure 5a-d illustrates for the example site pair from Figure 4a,b how we tested the hypothesis. We calculated the mean gamma phase relation over trials (weighted by the power per trial) and named it the “good” phase relation (indicated by the yellow line in Fig. 5a,c). RTs on a given trial should be predicted by the degree of deviation from this good phase relation. We rotated all phase relations by a fixed phase, such that the mean phase relation was at zero (Fig. 5a was rotated into Fig. 5c). We binned trials according to their phase relation and averaged RTs per bin (Fig. 5b). After rotation (Fig. 5b was rotated into Fig. 5d), this revealed that phase relations close to the mean indeed resulted in particularly short RTs. We applied the same analysis steps to all interareal site pairs (Fig. 5e), and this confirmed the effect in the population of site pairs.

**Figure 5.**
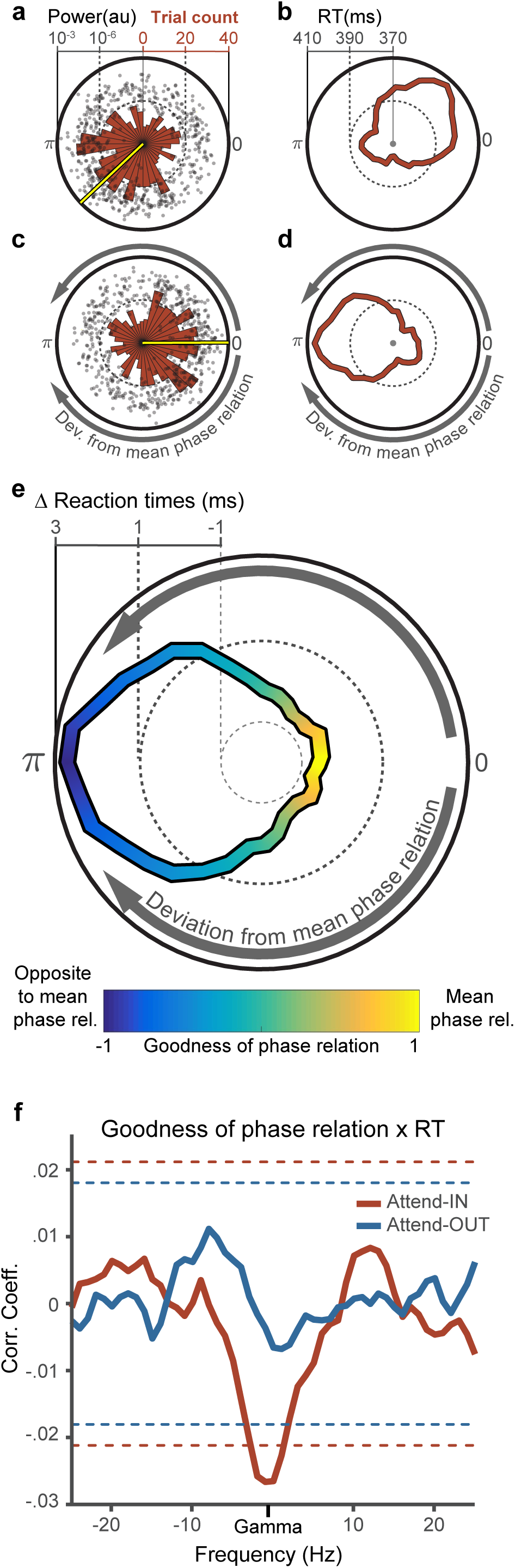
Trial-by-trial deviation from mean interareal gamma-band phase relation predicts reaction time. Panels (a-d) illustrate, using the example pair of Fig. 4a during attend-IN, that all phase relations of a given pair were rotated such that the mean phase relation was at zero. This rotation was applied to allow combination over site pairs, and to investigate whether RTs depended systematically on the deviation from the mean phase relation. (a) Each dot represents the gamma phase relation between V1 and V4 in one trial. The polar histogram in red shows the corresponding distribution. The yellow bar represents the mean gamma phase relation, corresponding to the phase relation at which gamma-band synchronization occurred. (b) Polar plot showing reaction time as a function of the V1-V4 gamma phase relation (copy of Fig. 4a). (c,d) Same as a,b, after rotating all phase relations such that the mean phase relation is at zero. Note that the rotation is based on the mean phase relation shown in a, and it reveals that RTs are not just correlated with phase relation, but systematically decrease with decreasing deviation from the mean phase relation. (e) Same as d, but average for all V1-V4 site pairs (after demeaning RTs). Color code reflects the cosine of the deviation from the mean phase relation, referred to as “goodness of phase relation” (GPR). (f) Spectrum of correlation coefficients between the GPR and RT, separately for attend-IN (red) and attend-OUT (blue). The dashed lines indicate the significance thresholds (p<0.05) obtained with a non-parametric randomization test including correction for multiple comparisons.

To quantify the observed relation between the deviation from the mean phase relation and RT, we took for each trial the cosine of the phase-relation deviation as a metric of that trial’s “goodness of phase relation” (GPR), such that trials with a good (bad) gamma phase relation obtained values close to one (minus one), as indicated by the color scale in Figure 5e. A linear correlation analysis, based on single-trial GPR and RT values without binning, revealed that good V1-V4 gamma phase relations resulted in short RTs (Fig. 5f; p<0.05, non-parametric randomization test corrected for multiple comparisons across frequencies). This effect was present during the attend-IN condition (red line), but not during the attend-OUT condition (blue line).

Note that the modulation of RTs by the GPRs is larger in the example (Fig. 5d) than in the average (Fig. 5e). We found that the correlation between GPRs and RTs was itself correlated with the strength of V1-V4 PPC (r=-0.225, p<0.004; calculated at gamma peak frequency for attend-IN; negative value reflects the negative GPRxRT correlation). This suggests that the trial-by-trial gamma phase relations of the interareal site pairs showing strong coherence for a given stimulus have a strong influence on the communication of that stimulus and thereby explain a substantial fraction of RT variability; The gamma phase relations of pairs with weaker coherence contribute less to communication or potentially show less coherence and RT modulation due to lower signal-to-noise recordings.

### RT modulation by interareal phase-relation is not explained by effects of local power or phase

Finally, we investigated to which degree the RT-predictive effect is specific to the V1-V4 gamma phase relation, rather than the gamma power in V1 or V4. Previous studies have shown that the gamma rhythm in V1 entrains the gamma rhythm in V4 in a feedforward manner (Bastos et al., 2015b; Bosman et al., 2012; Michalareas et al., 2016; van Kerkoerle et al., 2014). Thus, good gamma phase relations between V1 and V4 might be driven by strong gamma-band activity inside V1, and thereby the RT-predictive effect of the interareal phase relation might reduce to an effect of V1 (and/or V4) power. To test this, we first performed a linear regression with RT as dependent variable and the GPR as independent variable (Supplementary Fig. S2a). We then included the power in V1 and V4 as additional independent variables, and this left the results qualitatively unchanged (Supplementary Fig. S2b). Lastly, we repeated this control analysis, but adding as independent variable not the frequency-wise power but the power value at its area-wise peak frequency, which again did not change results (Supplementary Fig. S2c). Note that we replicate prior results showing that gamma power in V4 is negatively correlated with RT during attend-IN (Supplementary Fig. S3b) (Womelsdorf et al., 2006), and we furthermore find that gamma power in V1 is negatively correlated with RT during both attend-IN and attend-OUT (Supplementary Fig. S3a).

The control analysis using multiple linear regression relies on the assumption that the relations between RT and the different neuronal metrics are linear. We therefore performed an additional analysis that avoids this assumption by using stratification for gamma power. First, we split trials in two equal halves based on the median deviation from the mean gamma phase relation. We refer to these halves as “phase-relation conditions”. Then we sorted trials, separately for each phase-relation condition, according to gamma power, into nine bins, stratified conditions per bin for gamma power, and compared RTs between conditions. This revealed that RTs were significantly shorter for good than bad gamma phase relations, even after eliminating potential influences of power. This held after both, stratification for gamma power in V1 (p<0.001), and V4 (p<0.001; non-parametric randomization test).

The stratification approach allowed us to apply the same control for the gamma phase in V1 or V4 at the moment, when the stimulus change begins. A previous study had shown that the gamma phase in V4 around the time of a stimulus change can be predictive of reaction times (Ni et al., 2016). We did not find any significant correlation between either the V1 or the V4 phase and the V1-V4 phase relation, for any one of the investigated frequencies. Nevertheless, to test whether eventual effects of the local phases in V1 or V4 explain the effect of the V1-V4 phase relation, we applied phase stratification. We used the same phase-relation conditions as for power stratification. Then, as described above for gamma power, we stratified for gamma phase and compared RTs between conditions. This again confirmed shorter RTs for good gamma phase relations after both, stratifying for V1 gamma phase (p<0.001) and V4 gamma phase (p<0.001).

## DISCUSSION

In summary, we have shown that gamma synchronization between V1 and V4 improves behavioral performance. When trials were median split by their reaction times, faster RTs were preceded by stronger interareal synchronization in the gamma band. In fact, the trial-by-trial V1-V4 phase relation in the gamma band just before the behaviorally relevant stimulus change correlated with the behavioral RTs. Most importantly, the mean interareal phase relation, i.e. the phase relation at which synchronization occurs, resulted in the shortest RTs. When the phase relation deviated from the mean, this systematically slowed behavioral RTs. This shows that the gamma phase relation, at which V1 and V4 synchronize, is optimal for stimulus transmission to motor output. These effects occurred only, when the investigated gamma-band synchronization was induced by the attended stimulus, i.e. the stimulus that also triggered the behavioral response. This demonstrates that RTs depend specifically on the interareal synchronization involved in the transmission of the behaviorally relevant stimulus, rather than on fluctuations in overall arousal. Our results show that interareal communication depends on interareal coherence, directly supporting the central prediction of the CTC hypothesis.

Granger causality (GC) analyses have shown that the gamma rhythm in V1 entrains the gamma in V4 much more than vice versa (Bosman et al., 2012; Richter et al., 2017; van Kerkoerle et al., 2014). Across many pairs of visual areas, GC analyses show that interareal gamma entrainment generally proceeds along anatomical feedforward projections (Bastos et al., 2015b; Michalareas et al., 2016). Additionally, the available literature suggests that gamma-band synchronization among early visual areas V1, V2 and V4 is linked to the patterns of interareal monosynaptic connections (Bastos et al., 2015b; Roberts et al., 2013; Zandvakili and Kohn, 2015). Thus, the V1-V4 gamma phase relation is partly dependent on the strength of the driving local gamma-band rhythm in V1. Therefore, we investigated whether the relation between V1-V4 synchronization and RTs can be explained by local activity inside V1 or V4. Control analyses that stratified for local activities left the effect of interareal synchronization on behavior largely unchanged. As V1-V4 gamma synchronization is likely due to V1 gamma entraining V4 gamma through the respective monosynaptic feedforward projections, any explanation that invokes additional areas as common drivers is highly unlikely.

Several previous studies have shown that RTs can be predicted by local neuronal activity in individual visual areas. RTs in response to behaviorally relevant events can be predicted by neuronal firing rates evoked by these events in primary (Lee et al., 2010) and higher (Galashan et al., 2013; Womelsdorf et al., 2006) visual areas. Furthermore, RTs can be predicted by the strength and the absolute phase of local stimulus-induced gamma-band activity in visual areas of humans (Hoogenboom et al., 2010) and non-human primates (Ni et al., 2016; Womelsdorf et al., 2006). At the same time, numerous studies have shown that many cognitive functions, including perceptual organization, attention, working memory, and long-term memory encoding, do not only rely on local neuronal activity, but often specifically on interareal synchronization (Bosman et al., 2012; Buschman and Miller, 2007; Fell et al., 2001; Gregoriou et al., 2009; Grothe et al., 2012; Liebe et al., 2012; Salazar et al., 2012; van Kerkoerle et al., 2014). These findings have inspired the CTC hypothesis, stating that efficient interareal communication depends on interareal synchronization. Many recent studies have confirmed different predictions of the CTC hypothesis, such as the rhythmic gain modulation (Ni et al., 2016), and the selective interareal synchronization for attended information (Bosman et al., 2012; Grothe et al., 2012). The current study shows directly that an interareal phase relation that is close to the mean phase relation is optimal for information transmission as reflected in short RTs.

We found that behavioral response times are only predicted by interareal gamma phase relations, when gamma rhythms are induced by the attended, i.e. behaviorally relevant, stimulus. This supports the hypothesis that attentional selection is implemented by increased interareal communication (Fries, 2015). This notion has been supported by several studies using numerous different approaches. Firing rate recordings in higher visual areas combined with modeling have revealed that attentional effects on firing rates can be explained if attention enhances interareal gain (Reynolds et al., 1999; Ruff and Cohen, 2017). This was confirmed experimentally, when gain was directly quantified by the response of a higher area to the electrical microstimulation of a lower area. Selective attention enhances both the gain of LGN input to V1 (Briggs et al., 2013) and the gain of V1 input to MT (Ruff and Cohen, 2017). Finally, coherence and Granger causality between neuronal groups in different areas is enhanced, when they process attended stimuli (Bosman et al., 2012; Gregoriou et al., 2012; Gregoriou et al., 2009; Grothe et al., 2012; Richter et al., 2017; Saalmann et al., 2012; Zhou et al., 2016).

An interesting avenue for future research will be to investigate further interareal links on the way from V1 to motor cortex. The dependence of RTs on interareal synchronization that we demonstrate here for the V1-V4 link might hold along the entire way, or it might be transformed into an effect on firing rates or a synchronization in different frequency bands. Another important task for future studies will be to provide further evidence for a causal relevance of the observed relationship. Optogenetics affords the opportunity to generate gamma band activity in visual areas in-vivo (Ni et al., 2016), and to modulate the phase and frequency of induced gamma (Akam et al., 2012). With these tools, it should be possible to generate interareal synchronization at pre-specified phase relations. The results presented here directly lead to the prediction that optogenetically controlled interareal phase relations should modulate the efficiency of interareal communication and ultimately affect behavior.

## METHODS

### Electrophysiological Recording and Signal Processing

All experimental procedures were approved by the ethics committee of Radboud University Nijmegen (Nijmegen, The Netherlands). Data from two adult male macaque monkeys (macaca mulatta) were collected for this study. Parts of the data have been used in other publications (Bastos et al., 2015a; Bastos et al., 2015b; Bosman et al., 2012; Brunet et al., 2015; Brunet et al., 2014a; Brunet et al., 2014b; Hindriks et al., 2017; Lewis et al., 2016a; Lewis et al., 2016b; Pinotsis et al., 2014; Richter et al., 2015, 2017; Rubehn et al., 2009; Spyropoulos et al., 2017; Vinck et al., 2015).

LFP recordings were made via a 252 channel electrocorticographic grid (ECoG) subdurally implanted over the left hemisphere (Bastos et al., 2015b; Rubehn et al., 2009). Signals were filtered with a passband of 0.159 8000 Hz and sampled at approximately 32 kHz (Neuralynx Digital Lynx system). Offline, signals were low-pass filtered at 250 Hz and downsampled to 1 kHz. The electrodes were distributed over eight 32-channel headstages, and referenced against a silver wire implanted onto the dura overlying the opposite hemisphere. The electrodes were re-referenced via a bipolar scheme to improve signal localization, cancel the common reference, and reject noise specific to the headstage. The bipolar derivation scheme subtracted the recordings from neighboring electrodes (spaced 2.5 mm) that shared a headstage, resulting in 218 bipolar derivations, referred to as “recording sites” or just “sites” (see (Bastos et al., 2015b) for a detailed description of the re-referencing procedure). Areas V1 and V4 used here were defined based on comparison of the electrode locations, coregistered to each monkey’s anatomical MRI and warped to the F99 template brain in CARET (Van Essen, 2012), with multiple cortical atlases of the macaque (see (Bastos et al., 2015b) for a detailed description). Recording sites were coregistered to a common template (INIA19(Rohlfing et al., 2012)), as were the area definitions based on multiple cortical atlases. Based on these area definitions, 77 recording sites were selected from area V1 (monkey K: 29; monkey P: 48), 31 from area V4 (monkey K: 17; monkey P: 14). All analyses performed in this study were done on the sites that showed highest (top 1/3) visually induced gamma-band activity within areas V1 and V4 (Fig. 2a,b), resulting in a subset of 26 recording sites from area V1 (monkey K: 10; monkey P: 16), and 11 from area V4 (monkey K: 6; monkey P: 5).

All signal processing was conducted in MATLAB (Math-Works, USA) using the FieldTrip toolbox (http://www.fieldtriptoolbox.org/) (Oostenveld et al., 2011). Line noise was removed by subtracting the 50, 100, and 150 Hz components estimated through a discrete Fourier transform. Trial epochs for each site were demeaned by subtracting the mean over all time points in the epoch. Epochs with any site having a variance of greater than 5 times the variance based on all data from that same site in the same session were rejected. In addition, epochs were manually inspected, and epochs with artifacts were rejected. Subsequently, per recording site, the signals from all remaining epochs of a given session were divided by the standard deviation of the signal across all those remaining epochs, and the resulting z transformed signals were combined across sessions.

### Stimuli and Task

Stimuli were presented on a CRT monitor (120 Hz non-interlaced) in a dimly lit booth and controlled by CORTEX software (https://www.nimh.nih.gov/labs-at-nimh/research-areas/clinics-and-labs/ln/shn/software-projects.shtml). The paradigm is illustrated in Figure 1. Upon touching a bar, a fixation point was presented, and the monkey’s gaze was required to remain inside the fixation window throughout the trial (monkey K: 0.85 deg radius, monkey P: 1 deg radius). Otherwise the trial would be terminated and a new trial initiated. Upon the acquisition of central fixation, and after an 0.8 s pre-stimulus interval had elapsed, two isoluminant and isoeccentric drifting sinusoidal gratings were presented, one in each visual hemifield (diameter: 3 deg, spatial frequency: ≈1 cycle/deg, drift velocity: ≈1 deg/s, resulting temporal frequency: ≈1 cycle/s, contrast: 100%). Blue and yellow tints were randomly assigned to each of the gratings on each trial. Following a random delay interval (monkey K: 1 - 1.5 s; monkey P: 0.8 - 1.3 s), the color of the central fixation point changed to match one of the drifting gratings, which indicated that the matching grating was the behaviorally relevant or “target” stimulus. Thus, fixation point color acted as the attentional cue. When attention was directed to the stimulus in the visual hemifield contralateral (ipsilateral) to the recorded left hemisphere, this is addressed as the attend-IN (attend-OUT) condition. In each trial, two times were randomly drawn according to a slowly rising hazard rate to fall into an interval of 0.75 5 s (monkey K), and 0.75 4 s (monkey P) after stimulus onset. These two times were randomly assigned to be the change times of the target and the distracter. Bar releases 0.15 - 0.5 s after target changes were rewarded. Overall, 94% and 84% of all target changes were correctly reported by monkey K and monkey P, respectively (excluding fixation breaks). On half of the trials, the target changed first, and a corresponding behavioral response terminated the trial. Only those trials are analyzed here. The stimulus changes were subtle changes in shape consisting of a transient bending of the bars of the respective grating (0.15 s duration of the full bending cycle). All analyses presented here were performed on a window of 200 ms immediately preceding stimulus change (unless specified in text). We excluded trials in which the attention cue was presented less than one second before the target change, to ensure that cue processing was fully completed and attention deployed by the time of the target change. A total of 2550 trials (monkey K: 1149; monkey P: 1401) were used in this study.

### Spectral analysis

Each 200 ms epoch was multiplied with a Hann taper, zero padded to 1 s, and Fourier transformed, resulting in an FFT spectrum with a frequency resolution of 1 Hz. In the following, we will denote the arrays of Fourier spectra for all *K* epochs (*k* = 1,…, *K*) from the *i*-th site in V1 and the *j*-th site in V4 as *F*_*ik*_(*f*) and *F*_*jk*_(*f*), respectively, with *f* being the frequency in Hz. Spectral power was derived as the squared magnitude of the complex Fourier spectra. The relative power change shown in Figure 2c-f was computed as percent change relative to a baseline period of 200 ms before stimulus onset. Cross-spectral densities (CSDs) were estimated for each pair *ij* of V1 and V4 recording sites, for each frequency *f*, and each epoch *k* as

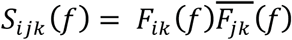

where 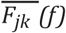 is the conjugate Fourier spectrum for V4. The magnitude of the CSD, |*S*_*ijk*_ (*f*)| reflects the product of spectral energies in V1 and V4, and the argument (angle) of the CSD reflects the phase relations between V1 and V4.

Phase locking was quantified by deriving from the CSD the pairwise phase consistency (PPC) metric, a phase locking metric that is not biased by the number of epochs (Vinck et al., 2010b).

### Phase Relation Analysis

We aimed at quantifying whether behavioral RTs were related on a trial-by-trial basis to the V1-V4 phase relation. We hypothesized that the mean V1-V4 phase relation is optimal for interareal communication and thereby leads to the shortest RTs. Note that the raw phase relation between a given V1-V4 site pair cannot be directly interpreted, because the absolute LFP phase depends on numerous accidental factors like the geometric relationship between source and electrode, and additionally the bipolar derivation used to remove the common recording reference incurs arbitrary phase rotations. Yet, irrespective of this, the mean phase relation reflects the phase relation of synchronization, and the phase relation in each trial can be expressed in terms of its deviation from this mean phase relation. Therefore, for each interareal site pair and epoch separately, we first calculated the mean phase relation and subtracted it from each epoch’s phase relations. The mean phase relation was determined by computing the mean resultant vector across epochs, defined as

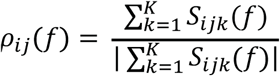

For each epoch, we then determined the phase deviation from the mean phase relation by computing the “rotated CSD”, by multiplying with the conjugate of the mean resultant vector,

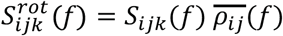

The argument (angle) of the rotated CSD, 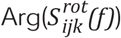, then corresponds to the phase difference between the CSD in the k-th trial and the mean phase relation. According to our hypothesis, any deviation from the average phase relation results in sub-optimal interareal communication. After rotating the average phase relation into zero, we could quantify the deviation of an epoch’s phase relation from the mean phase relation by taking the cosine of the angle of the rotated CSD. We refer to this metric as the goodness of phase relation (*GPR*):

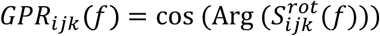

According to the hypothesis, a *GPR* value of 1 corresponds to a good phase relation, and a *GPR* value of −1 corresponds to a bad phase relation. Note that the *GPR* ignores the direction, in which a given epoch’s phase relation deviates from the average (see Fig. 5e).

### Statistical inference

All statistical inferences were based on the combined data from both animals, resulting in a fixed-effect analysis for our sample. All significance thresholds were computed using non-parametric permutation statistics (Maris and Oostenveld, 2007). For the median split analysis, the observed PPC spectra (without randomization) were derived by 1) calculating PPC spectra across all epochs of a given condition in a given animal, separately for all V1-V4 site pairs, 2) averaging PPC spectra across those site pairs, and 3) averaging across the two animals after aligning the peaks of the respective frequency bands. Surrogate distributions were then generated by randomly distributing epochs in two conditions, maintaining the sample sizes of the original conditions (short RTs versus long RTs) (Maris et al., 2007). This procedure was repeated 1000 times to produce a randomized null distribution. Then, the same steps as for the observed PPC spectra were followed. For each randomization, the maximal absolute difference value across all frequencies was retained and placed into the randomization distribution. The observed differences were compared against the distribution of maximal absolute differences. This procedure corrects for multiple comparisons across frequencies. A similar approach was used in the phase relation analysis. A surrogate distribution was generated under the hypothesis that there is no correlation between RTs and the preceding phase relation between V1 and V4. So, in each randomization, the correlation coefficients were calculated on randomly shuffled RTs. The observed coefficients were then compared against the surrogate distribution of maximal absolute coefficients.

## ACKNOWLEDGEMENTS

We thank Craig Richter for preprocessing the data, Julien Vezoli for help with assigning electrode and recording sites to brain areas, and Martin Vinck for comments on the analysis description. PF acknowledges grant support by DFG (SPP 1665, FOR 1847, FR2557/5-1-CORNET, FR2557/6-1-NeuroTMR), EU (HEALTH-F2-2008-200728BrainSynch, FP7-604102-HBP, FP7-600730-Magnetrodes), a European Young Investigator Award, NIH (1U54MH091657-WU-Minn-Consortium-HCP), and LOEWE (NeFF).

## AUTHOR CONTRIBUTIONS

C.A.B. and P.F. designed the experiments; C.A.B. collected the data; G.R. and P.F. analyzed the data; G.R. and P.F. wrote the paper.

## COMPETING FINANCIAL INTERESTS

The authors declare no competing financial interests.

---

**Figure S1.**
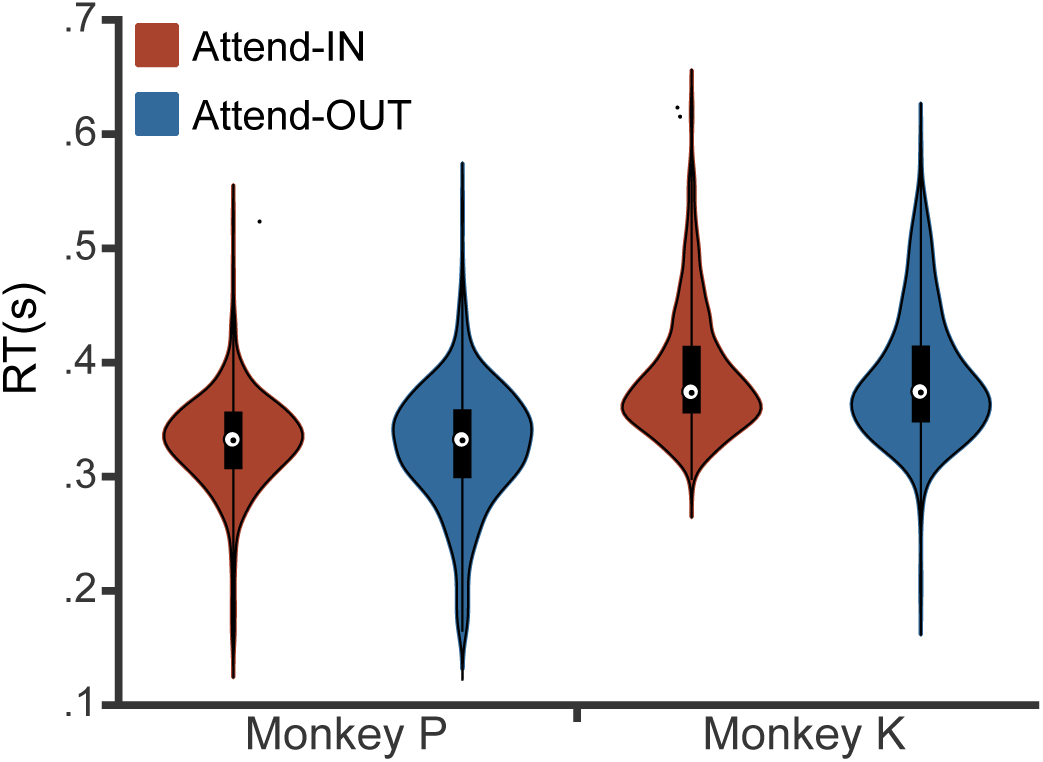
Distributions of reaction times, separately for attend-IN and attend-OUT and for each of the two monkeys. RT distributions are shown as color-filled outlines (“violin-plots”), whose width reflects the relative number of trials with the respective RT. The violin-plots contain inside a box plot, i.e. a white dot reflecting the median, a rectangle with its upper and lower edges reflecting the upper and lower quartile, and whiskers reflecting 3 times the interquartile range. The individual dots outside the violin plots reflects RTs outside the whisker range.

**Figure S2.**
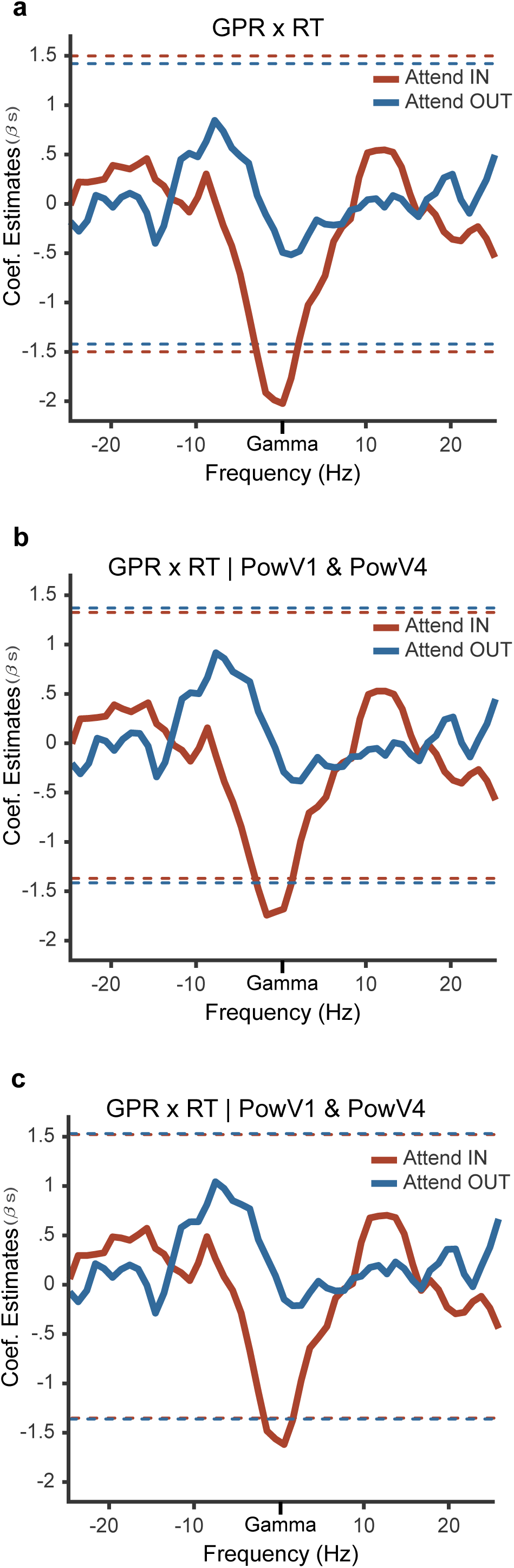
Control for effects of local power using multiple linear regression. (a) Spectrum of correlation coefficients between the GPR and RT, separately for attend-IN (red) and attend-OUT (blue). The dashed lines indicate the significance thresholds (p<0.05) obtained with a non-parametric randomization test including correction for multiple comparisons. (b) Same as a but including power in V1 and V4 as independent variables. (c) Same as b but using power at the gamma peak frequency of the respective area.

**Figure S3.**
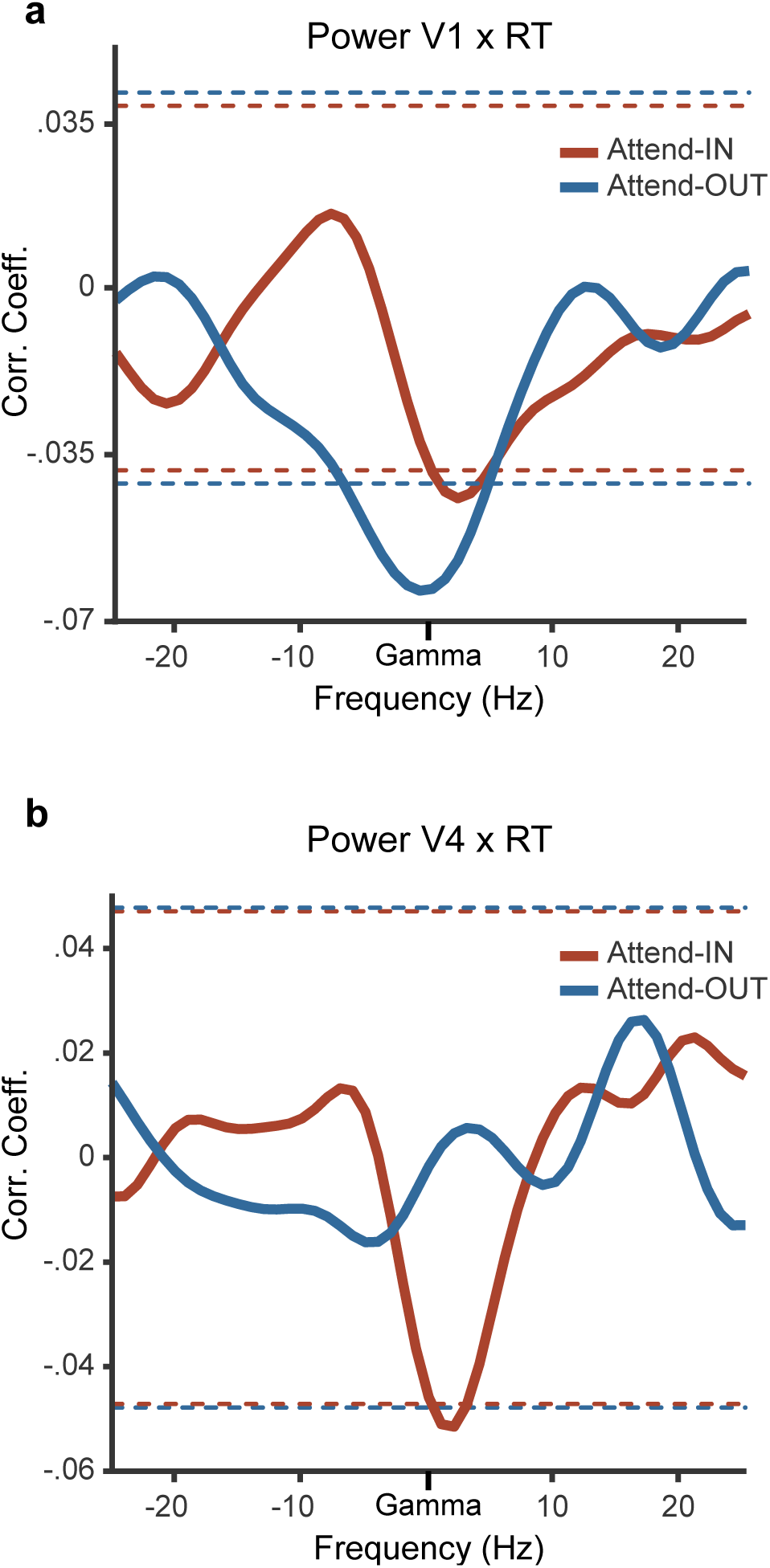
Relation between local power and reaction times. (a) Pearson correlation coefficient between power in V1 and RT, as a function of frequency after alignment to the individual gamma power peak frequency. (b) Same as a, but for area V4. Data from attend-IN (attend-OUT) are shown in red (blue). The dashed lines indicate the significance thresholds (p<0.05) obtained with a non-parametric randomization test including correction for multiple comparisons.

